# Developing a zebrafish xenograft model of diffuse midline glioma

**DOI:** 10.1101/2025.03.31.646163

**Authors:** Kaixuan Wang, Gianna Graziano, Anneliese Ceisel, Huanhuan Xiao, Shreya Banerjee, Yuran Yu, Marina Venero Galanternik, Brant M Weinstein, Charles G. Eberhart, Jeff Mumm, Eric Raabe

## Abstract

Diffuse midline glioma (DMG) is a highly aggressive brain tumor that predominantly affects children. Conventional treatments such as radiation therapy can control progression for a time, but DMG kills nearly 100 percent of patients. Although murine models have provided critical insights into the biology of DMG and in assessing new therapeutic strategies, they are not suitable for high-throughput screening to identify and profile novel therapies due to technical challenges, ethical considerations and high cost. Zebrafish (*Danio rerio*) is an established vertebrate model for large-scale drug screening, and zebrafish have demonstrated the ability to replicate the key biological and pathlogical aspects of human malignancies.

Here, we developed a novel method for transplanting human DMG cells into large numbers of zebrafish embyros to speed the assessment of anti-tumor drug efficacy *in vivo* and thereby facilitate the development of novel therapeutics for clinical translation. We transplanted red fluorescent protein (RFP)-labeled, patient-derived DMG cell lines into zebrafish blastulas. Remarkably, many DMG cells migrate into the developing brain and are present in the midline of the brain 24 hours after blastula injection. Tumor cell burden was monitored by measuring RFP fluorescence intensity changes over time. Time-course images of transplanted tumor cell volumes were acquired, and the interactions between transplanted DMG cells and microglial cells were further analyzed using Imaris software. We have developed a simple and rapid transplantation protocol to establish a zebrafish xenograft model of DMG. Our method involves transplanting DMG cells into the blastula stage (1000 cell stage) of zebrafish embryos, which does not require complex surgical techniques. This approach allows for the transplantation of hundreds of embryos per hour, significantly increasing the efficiency of creating DMG zebrafish xenografts that are suitable for high-throughput drug and gene discovery screens.

## Introduction

Diffuse midline gliomas (DMGs) are rare central nervous system (CNS) tumors that originate in midline structures, including the thalamus, brainstem, and spinal cord.[1,2] Classified as high-grade gliomas, DMGs are aggressive malignant tumors with a poor prognosis. They primarily affect children and young adults, with peak incidence observed in children aged 5 to 10 years.[3] [4] DMGs have a median survival of less than one year from diagnosis.[5] DMGs are highly metastatic tumors that infiltrate surrounding brain tissue, rendering surgical removal nearly impossible.[6,7] Additionally, DMGs exhibit resistance to traditional chemotherapy and radiation therapy, which are standard treatments for other types of brain tumors.[8] Given the challenges in treating DMGs, understanding their complex biology within a natural physiological environment and conducting high-throughput preclinical durg testing are essential. Enhancing the current animal models is a critical step toward achieving this goal.[9]

Xenograft models involve transplanting DMG cells or fresh brain tumor spheroids from patients into immunodeficient rodents. These models effectively replicate the genetic diversity and phenotypic characteristics of the original tumors.[10] Orthotopic xenografts of DMG in rodents, while valuable, face significant technical challenges and do not perfectly replicate the original tumor, and cannot conduct high-throughput drug screening. The diffusely infiltrative nature of these gliomas creates on average a 6 month latency between tumor implantation and the death of mice from tumor growth.[11] While this long period of tumor growth and infiltration phenocopies the primary human tumor, the delay in reaching endpoints such as survival or bioluminescent imaging of tumor growth increase the time and expense of pre-clinical experimentation.

In the last ten years, the zebrafish (Danio rerio) has become an important and clinically relevant vertebrate model for cancer research, aiding in the screening of small molecules that inhibit cancer and evaluating the toxicity of drugs.[12] [13] [14] [15] [16] [17] Zebrafish afford several key advantages, such as rapid development, small size, high reproductive capacity, and low-cost maintenance. Moreover, their lack of a functional adaptive immune system until 21 days post-fertilization makes them particularly well-suited for xenotransplantation experiments.[18] Furthermore, zebrafish are exceptionally well-suited for high-throughput drug or gene discovery screening and toxicological testing. They have already been utilized to develop adjuvant and metastatic therapies for uveal melanoma. [19] [17] Recent studies have underscored the outstanding capability of zebrafish to replicate disease features and their significant relevance in clinical settings. Utilizing transgenic models allows for the exploration of tumor cells within their microenvironment and the identification of key downstream pathways.[20] [16] However, intracranial transplantation of DMG cells into the brains of individual zebrafish larvae is technically demanding and time-consuming, currently limiting their use in high-throughput screens for potential therapies. A novel method that enables easy and rapid injection of DMG cells into the 1000-cell-stage zebrafish blastula could significantly streamline the process and save time. This approach could allow for the injection of hundreds of embryos per hour, facilitating high-throughput drug and gene discovery screens and accelerating clinical translation.

In this study, our objective is to establish a new zebrafish xenograft model that facilitates an easy and fast injection of DMG cells into zebrafish embryos. This method was adapted from established Glioblastoma (GBM) xenotransplantation protocols to optimize the engraftment and survival of DMG cells in the zebrafish model.[21] To begin, we collected zebrafish embryos at ~1000-cell stage for microinjection, using both wild-type and transgenic animals (Leptomeninges-YFP zebrafish and Microglia-YFP zebrafish). These models are generated by integrating transgenic DNA fragments into the zebrafish genome, enabling stable overexpression of fluorescent proteins in specific cell compartments, such as microglial cells, or leptomenigeal cells lining the ventricles. This approach facilitates visual confirmation of DMG localization and the investigation of interactions between DMG cells and microglia, providing valuable insights into the tumor microenvironment. At 3-4 days post injection (dpi), confocal microscopy was employed to monitor tumor progression and assess the interactions between DMG cells and leptomenigeal cells or microglial cells of the injected embryos. A fluorescence microplate reader assay was used to quantify the RFP signal intensity[22,23], providing insights into tumor cell proliferation and treatment responses (Figure 1). This approach has been used in other disease contexts for large-scale drug and gene screening in zebrafish larvae.[24–27]

**Figure 1:**
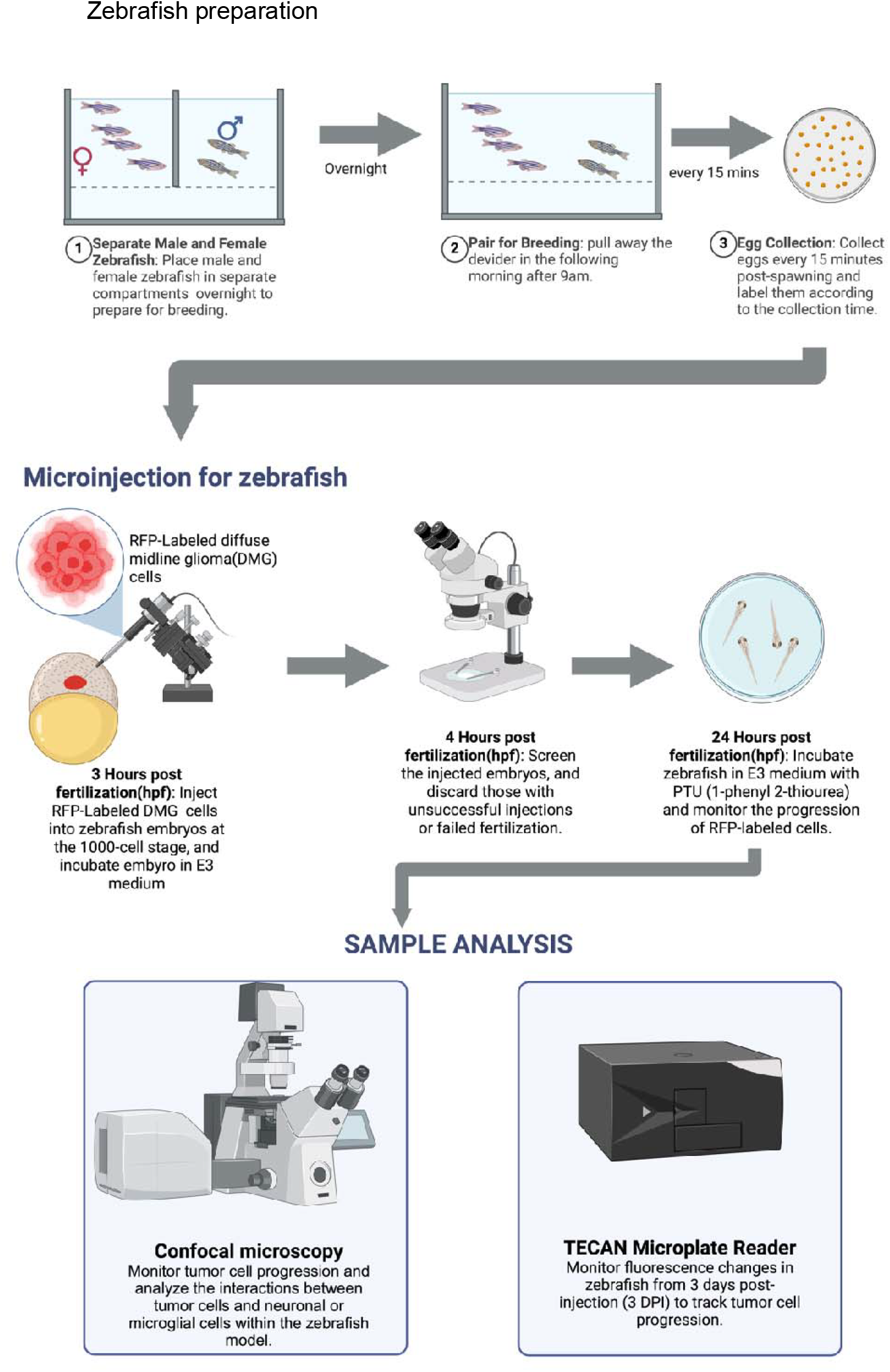
Illustrations of the zebrafish embryo preparation, microinjection, and sample analysis.

## 3. Materials

### 3.1 Cell lines and cell culture

Cell lines: JHH-DIPG-2JA, JHH-DIPG-17, HSJD-007

JHH-DIPG-2JA was developed from a post-treatment metastasis.

JHH-DIPG-17 was developed from a pre-treatment biopsy sample. HSJD-007 was developed from an autopsy sample at Hospital St. Joan de Deu in Barcelona, Spain and was provided by the kind gift of Angel Carcaboso. Cell lines were confirmed to harbor the H3K27M mutation and were validated by STR testing. DMG cells were cultured in DMEM/F12 DIPG proliferation medium (EF medium), prepared to a final composition of: 30% Ham’s F12 (Invitrogen, Cat# 11765-062), 70% DMEM (Invitrogen, Cat# 11965-118), 1% L-glutamine, 2% B27 supplement (Invitrogen, Cat# 12587-10), 20 ng/ml EGF (Peprotech, Cat# AF-100-15), 20 ng/ml FGF (Peprotech, Cat# 100-18B), and 5 µg/ml heparin.

### 3.2 Lentiviral transduction

For transduction, RFP-labeled lentivirus (Firefly Luciferase-T2A-RFP-IRES-Bsd Lentivirus, Biosettia, Cat# Glowcell-15b-1) was used. Blastcidin (Thermo Fisher Scientific, Cat# R21002), Polybrene (Millipore Sigma, Cat# H9268), and Accutase (Innovative Cell Technologies, Cat# A694-100ML) were also utilized in the transduction protocol.

### 3.3 Transgenic zebrafish

To create “Microglia-YFP” zebrafish, a macrophage-specific Gal4 driver line [*Tg(mpeg1:GAL4-VP16)gl24*] was crossed to a UAS reporter-effector line co-expressing YFP and a nitroreductase 2.0 (NTR 2.0) [*Tg(5xUAS:GAP-TagYFP-P2A-NfsB_Vv_F70A/F108Y)jh513*]. To create “Leptomeninges-YFP” zebrafish, a leptomeninges-specific Gal4 driver line [*Tg(epd:Gal4)*] was crossed to a different UAS reporter-effector line co-expressing YFP and nitroreductase 2.0 (NTR 2.0)[28] *Tg(5xUAS:GAP-TagYFP-P2A-NfsB_Vv_F70A/F108Y, he:tag-BFP2)jh552*

### 3.4 Animal Studies

All procedures were conducted in compliance with the guidelines of the Office of Laboratory Animal Welfare (OLAW) for zebrafish research and under an approved protocol by the Johns Hopkins University Animal Care and Use Committee. Zebrafish were maintained under standard laboratory conditions at 28.5°C with a 14-hour light and 10-hour dark cycle. The larvae designated for imaging were treated with 1-phenyl 2-thiourea (PTU) starting at 1 day post-fertilization (dpf) to inhibit pigmentation.

### 3.3 Equipment and Materials

#### The following equipment and materials were used in this study

Confocal Microscope: Olympus FV1000 equipped with 515 nm and 559 nm laser lines. Stacked confocal images were captured using an XLUMPLFL 20X W NA: 0.95 objective, with a step size of 5 µm.

Fluorescence Microscope: Motic AE31.

Fluorescence microplate reader (TECAN Infinite M1000 PRO; excitation 555 nm, bandwidth 5 nm; emission 585 nm, bandwidth 10 nm)

Microinjection System: NARISHIGE IM-200 Microinjector.

Needle Puller: Sutter Instrument Model P-97, configured with the following settings: Heat 580, Pull 70, Velocity 90, Time 110.

Glass Capillaries: 4-inch Borosilicate Glass, 1.2 mm OD (Cat# 1B120-4, WPI). Microloader Tips: Caliber: 2 × 96 (Cat# EPE-930001007).

Microplate: 96-well, polypropylene (PP), U-bottom, black (Greiner Bio-One, Cat# 650209).

### 3.5 Other reagents

Tricane methanesulfonate anesthetic (Fisher, NC0342409, final concentration: 0.612 nM)

Low melt agarose (Fisher, BP1360–100)

E3 media: Prepare 1 liter of 100X E3 medium by dissolving 29.22 g of NaCl, 1.27 g of KCl, 3.33 g of CaCl_2_, and 3.97 g of MgSO_4_ in water. Adjust the pH to 7.4 using 5 M NaOH. To obtain 1× E3 medium, dilute the 100× E3 stock solution with water. The final solute concentrations in 1× E3 medium are NaCl (5 mM), KCl (0.17 mM), CaCl_22_(0.3 mM), and MgSO_4_ (0.33 mM).

PTU pigmentation inhibition agent (1-Phenyl-2-thiourea, Fisher-P023725G, final concentration: 200 nM)

## 4. Methods

### 4.1 Lentivirus transduction

The lentivirus used contains a blasticidin resistance gene for selection. Prior to transduction, cell lines were tested for blasticidin tolerance to determine the optimal concentration. Cells were seeded in a 24-well plate, with each well containing the appropriate number of cells (e.g., 20,000 cells per well for HSJD007). Eleven blasticidin concentration groups were prepared, each treatment was performed in duplicate wells, with concentrations ranging from 0 µg/ml to 10 µg/ml. Blasticidin was added, and cells were monitored for one week. Cell health was assessed daily under a microscope, and cell viability was confirmed using CellTiter-Blue. The lowest blasticidin concentration that effectively killed all cells was determined for use in subsequent experiments.

Lentivirus (Firefly Luciferase-T2A-RFP-IRES-Bsd Lentivirus, Biosettia, Cat# Glowcell-15b-1) was added to the cells based on the cell density in each dish. Depending on the cell line, 5-10 µg/ml of polybrene (Millipore Sigma, Cat# H9268) was used. After 24 hours of incubation, cells were centrifuged, and the medium was replaced with fresh culture medium. Two days post-transduction, cells were cultured in medium with the predetermined concentration of blasticidin for one week, with regular medium changes. Transduction efficiency was quantified by determining the percentage of RFP-positive DMG cells using fluorescence microscopy. And, for patient-derived DMG cells, single-cell sorting is not recommended, as it may result in the loss of many cells due to their large size and damage during dissociation and sorting.

### 4.2 Cell Preparation

#### 4.2.1 Thawing DMG Cell Lines (JHH-DIPG-17, JHH-DIPG-2JA, HSJD-007)

Cryopreserved DMG cell lines were thawed by removing vials from liquid nitrogen storage and placing them directly into a 37°C water bath. Once thawed, cells were transferred gently using a 1 ml pipette, Pasteur pipette, or P1000 pipette tip into a 15 ml conical tube. Pre-warmed medium was added dropwise (~200 µl per drop) using a P1000 pipette, with gentle swirling after each drop to gradually equilibrate cells from their cryoprotectant solution to isotonic conditions. This process continued until a final volume of 4 ml was reached. Cells were then centrifuged, the supernatant was removed, and the cell pellet was resuspended in 10 ml of fresh DMEM/F12 DIPG proliferation medium (EF medium). Cells were seeded into T25 flasks with regular medium change.

#### 4.2.2 Cell Collection and Preparation for Zebrafish Embryo Injection

To prepare cells for injection into zebrafish embryos, the cell-containing medium was transferred to a 15 or 50 mL conical tube and centrifuged to collect the cell pellet. Given the tendency of DMG cells to form tumor spheroids, the supernatant was carefully aspirated, and Accutase was added to the cell pellet to promote dissociation into single cells. The suspension was then gently mixed and incubated in a 37°C water bath for 5 minutes. Following incubation, cells were centrifuged at 1000 g for 5 minutes. The Accutase was aspirated, and the cells were resuspended by pipetting with a 200 µl tip to ensure a single-cell suspension. The suspension was then passed through a 40 µm nylon mesh to further refine the single-cell preparation.[29]

Cell counts were performed using a hemocytometer under a fluorescence microscope to quantify both the total cell count and the proportion of RFP-labeled cells. The cell suspension was washed three times with ice-cold phenol red-free HBSS buffer to eliminate any residual growth factors or medium components that could be toxic to zebrafish embryos. Finally, cells were resuspended in ice-cold HBSS medium containing 2% polyvinylpyrrolidone (Sigma) to avoid clogging of the micro-injection capillary at a concentration of 100 cells/nl.[29]

### 4.3 Fish Preparation

Zebrafish breeding pairs were set up in the afternoon (the day before the injection) to ensure fertilized embryos were available on the injection day. The setup, illustrated in Figure 2A, includes the tank, breeding divider, inserted breeding ramp, and lid arranged from left to right. The tank was assembled with the divider positioned to separate male and female zebrafish.

**Figure 2:**
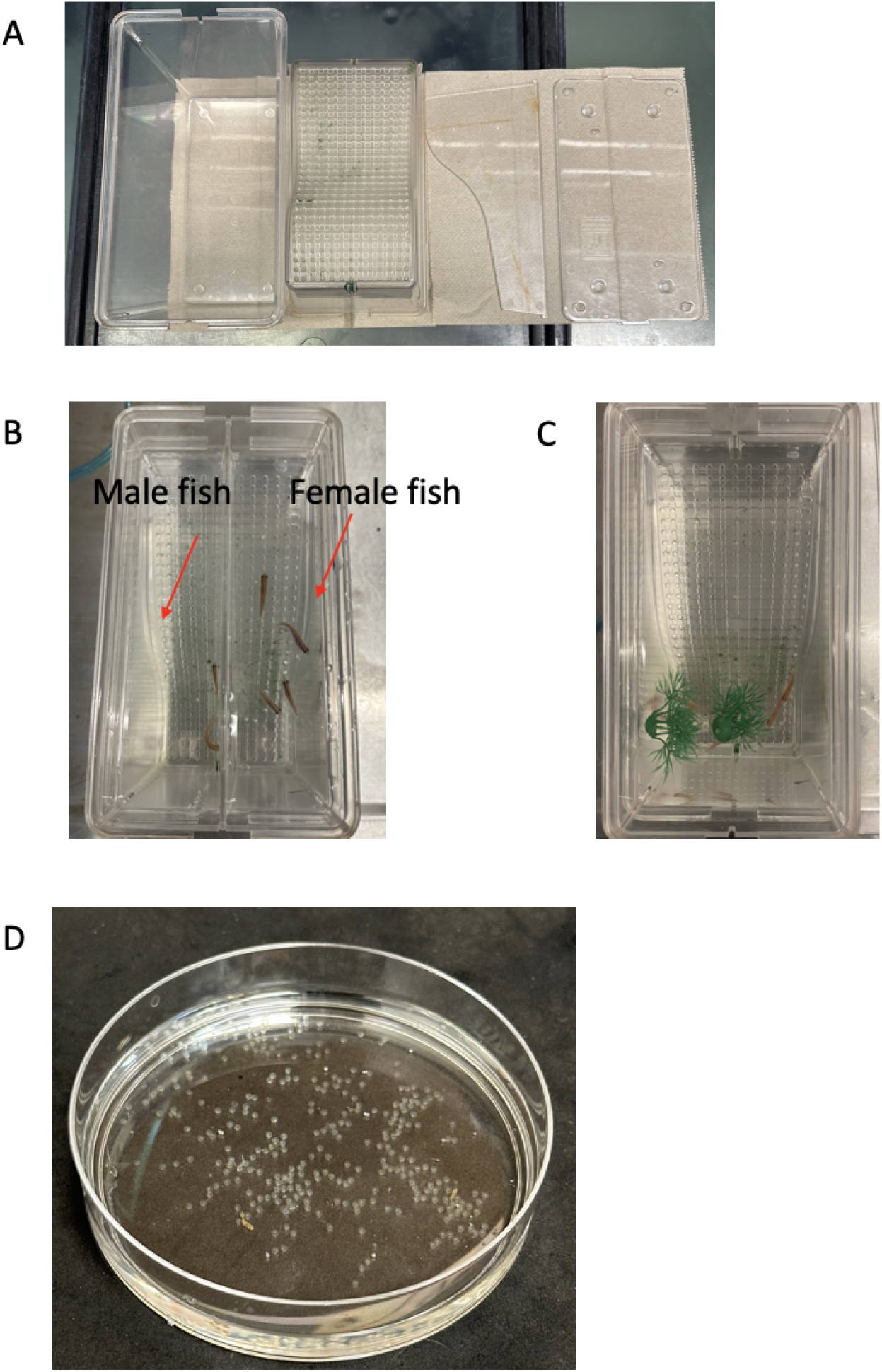
Breeding setup and procedure for zebrafish. **A)** Components of the breeding setup, including the tank, breeding ramp, dividers, and lid (from left to right). **B)** Zebrafish tank setup for overnight breeding. **C)** Breeding tank with the slide removed to initiate spawning. **D)** Collected embryos. Embryos are collected every 15 minutes and labeled with the corresponding collection time.

On the following morning, the divider was removed to allow breeding, and embryos were collected at 15-minute intervals. Each collection was labeled with the corresponding time of collection. The collected embryos were placed in a 28.5°C incubator. After 3 hours of incubation, the embryos were examined under a microscope to select ~1000-cell stage embryos for microinjection.

### 4.4 Fish Injection

When the zebrafish embryos were close to ~1000-cell stage, tumor cells were prepared for injection. Embryos were aligned in the mold (Figure 3C) and oriented using a pipette tip to such that the blastoderm was facing upwards, facilitating injection and minimizing injury (Figure 3D).

**Figure 3:**
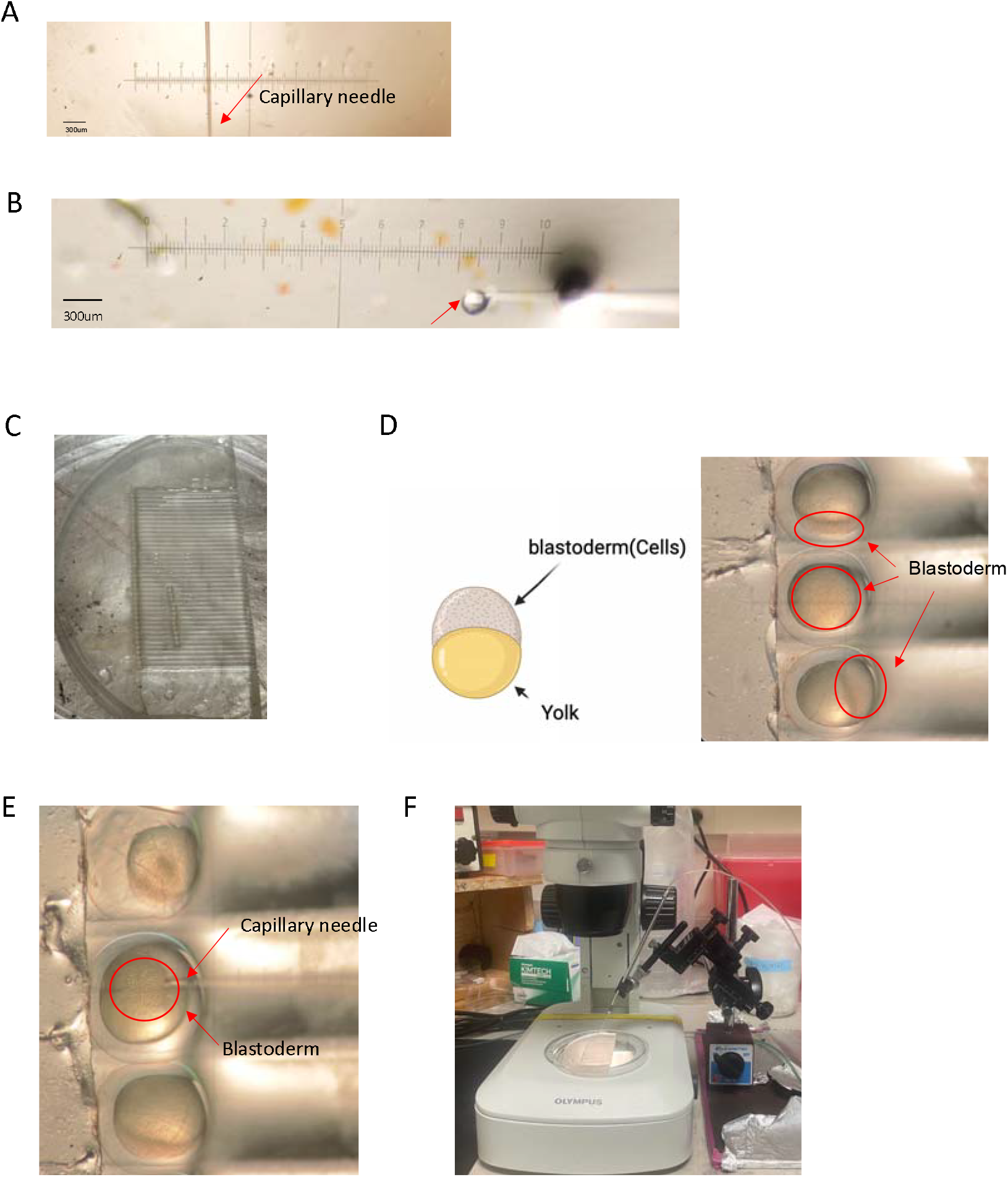
Microinjection procedure and apparatus for zebrafish embryos. **A)** Microinjection capillary used for injections, with ruler under the microscope. each tick mark represents 30 µm, with the needle trimmed to a diameter of approximately 20-25 µm. **B)** Measurement of the injection droplet size (with a diameter of approximately 130 µm and corresponding to a volume of approximately 1 nL) under the microscope to ensure accurate volume delivery. Approximately 100 tumor cells are injected into each embryo. **C)** Mold used to hold embryos during microinjection. **D)** The left cartoon illustrates the blastoderm and yolk of a zebrafish embryo at the 1000-cell stage. The right image shows orientation of a 1000-cell stage embryo for injection, with the blastoderm positioned upwards. **E)** The injection of embryos undergoing microinjection. **F)** Apparatus showing the microinjection holder and the needle angle (approximately 60 degrees).

#### Injection Setup

The air source and microinjector were activated, and the tips of injection needles were cut under a microscope as needed. Given the size of human cells (10~20 µm), the needles were trimmed using a sharp blade to a tip diameter of approximately 25–30 µm at a 45 angle, which facilitated smooth cell injection and minimized injury for the embryo. (Figure 3A). DMG cells were loaded into the needle capillary using a microloader (Cat# EPE-930001007), adding approximately 4 µl of cell suspension. The cells were kept on ice at all times to help prevent clogging within the needle capillary.

#### Injection Calibration

The cell suspension was prepared at 1 million cells per 10 µl to achieve the desired density (100 cells/nL). The needle was backfilled with DMG cells and securely mounted in the microinjector holder of the microinjection system. Injection solution was dispensed using a foot pedal. The needle tip was placed into mineral oil, and DMG cells were injected into the oil to produce droplets. The droplet size, with a diameter of approximately 130 µm and corresponding to a volume of approximately 1 nL, was adjusted and measured by fine-tuning the injection time and air pressure.(Figure 3B) After calibration, the capillary needle was positioned at a 60° angle for the injection procedure.

#### Embryo Injection Procedure

The embryos and needle were brought into focus under the microscope, ensuring that the injection angle remained at approximately 60° (Figure 3F). This angle helps prevent cell clumping of cells within the needle, minimizing the risk of flow obstruction and injection failures, such as empty injections or injections containing only buffer. Additionally, it reduces the likelihood of needle-induced injury to the embryo. During the experiment, mineral oil was used to check needle patency after each round of injections. If cell clumping was observed, the needle was replaced to ensure consistent performance.

#### Post-Injection Embryo Care

Following injection, embryos were transferred to E3 medium and immediately screened for the presence of RFP-labeled DMG cells under a fluorescence microscope. Positive embryos were selected and placed in new plates, ensuring adequate space to prevent overcrowding and subsequent embryo mortality. The morning after the injection, the E3 medium was replaced with E3 medium supplemented with 1-phenyl 2-thiourea (PTU) to inhibit pigmentation, thereby facilitating live imaging. Embyros were incubated in 28.5°C degree incubator.

Embryos were monitored daily, and dead embryos were removed promptly to prevent contamination. Fresh E3 + PTU medium was changed every other day. At five days post-injection, zebrafish were fed paramecia to facilate larval development.

### 4.5 Tracking Tumor Cell Progression in Zebrafish

Before the time-course imaging, the fish were anesthetized by transferring them into embryo medium supplemented with tricaine, allowing approximately 3 minutes for them to become unresponsive to touch. Subsequently, they were transferred to a transparent 96-well plate. A homemade hair loop or 10 µL pipette tip was employed to position and orient the fish appropriately. Time-course images were captured using fluorescence microscopy at the initial time point (0 hours), and subsequently at 24, 48, and 72 hours post-injection to monitor the migragation of tumor cells(Figure 5). After imaging, the fish were thoroughly washed to remove tricaine and placed back into E3 +PTU medium for incubating until the following imaging series.

The time-course images (Figure 4A, B) display an example of blastulas injected with JHH-DIPG-2JA cells or JHH-DIPG-17 cells. These images demonstrate the proliferation and migration of DMG cells inside the zebrafish.

**Figure 4:**
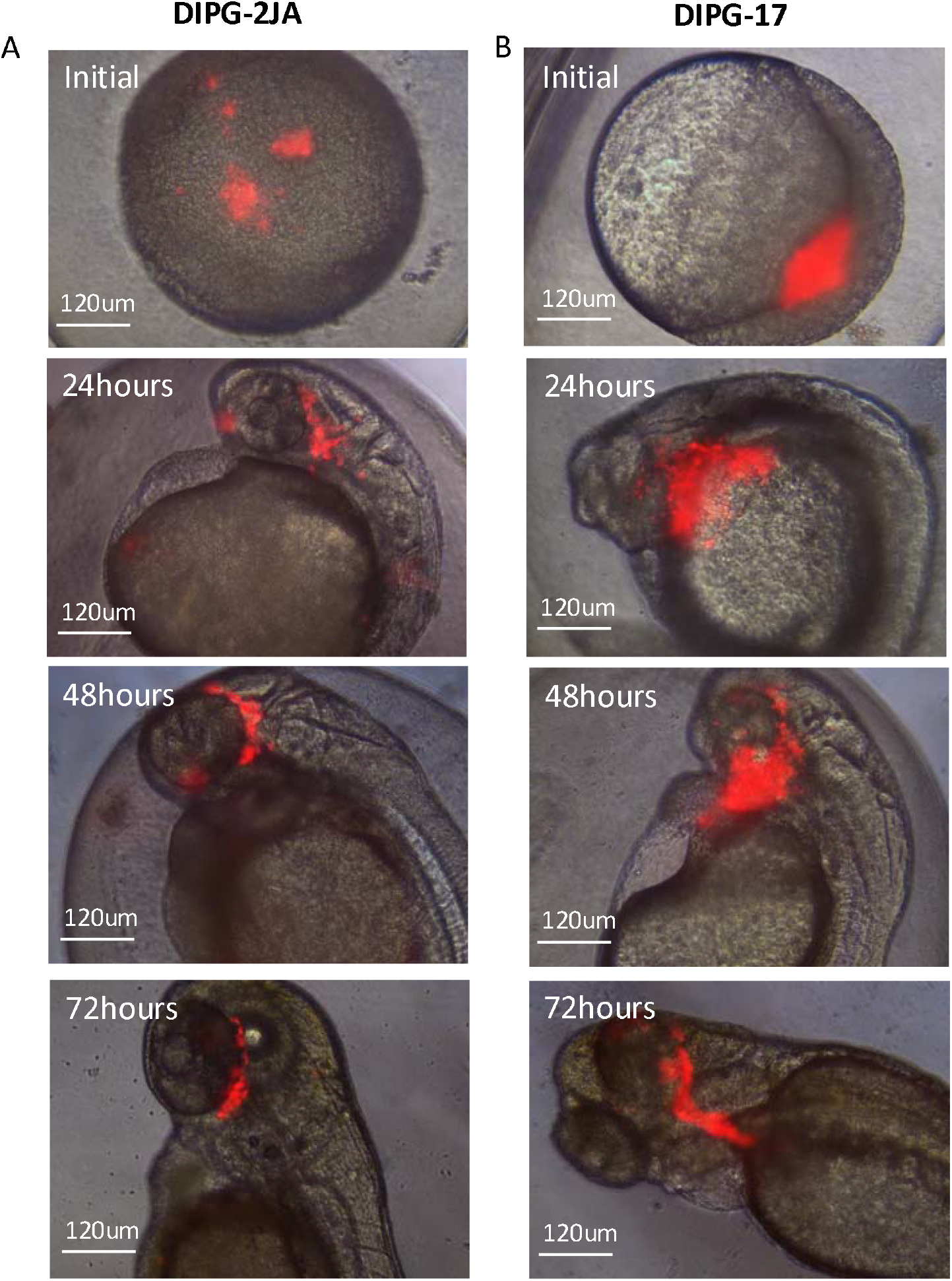
Time-course images after blastula injections. **A)** Time-course images showing the progression of DIPG-2JA cells following blastula-stage injection in zebrafish embryos. Fluorescent images were captured at the initial injection (0 hours), and at 24, 48, and 72 hours post-injection, illustrating the distribution of RFP-labeled tumor cells over time. Scale bar: 120 µm. **B)** Time-course images following blastula-stage injection of the DIPG-17 cell line in zebrafish embryos. Images were captured at the initial injection (0 hour) and at 24, 48, and 72 hours post-injection, showing the distribution of RFP-labeled DIPG-17 tumor cells over time. Scale bar: 120 µm.

### 4.6 Quantifying RFP-Labeled DMG Cell Fluorescence in Zebrafish

Fluorescence intensity was quantified using a TECAN microplate reader at 3 and/or 4dpi. Individual zebrafish were anesthetized with tricaine and placed in separate wells of a black 96-well plate (Figure 5A). Non-injected zebrafish were included for defining the fluorescent signal cutoff. To ensure accuracy and minimize variation in the microplate reader results, the zebrafish larvae were washed multiple times with clean PTU/E3 medium to remove any variations caused by impurities in the water.

**Figure 5.**
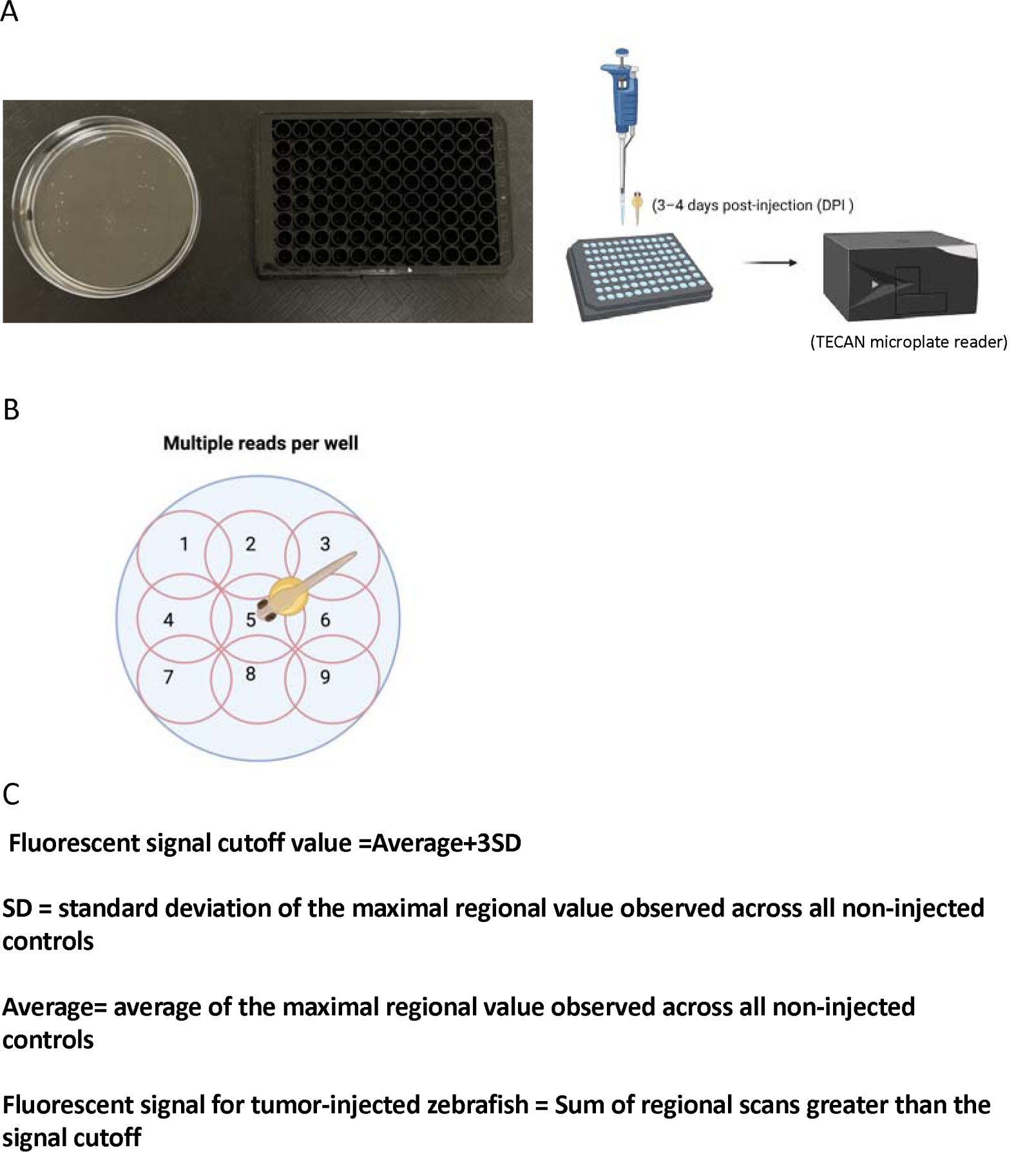
The setup and analysis of zebrafish fluorescence intensity by TECAN microplate reader. **A)** The left image shows a black U-shape 96-well plate and zebrafish larvae. The right image illustrates the procedure of placing zebrafish into the 96 wells followed by microplate reading. **B)** Alignment of the nine measurement points per well, representing the reading grid within the well. **C)** Non-injected zebrafish were used to establish a fluorescent signal cutoff value defined as the average of the maximal fluorescence intensity value for eachnon-injected zebrafish plus three standard deviations (avg + 3SD). Total fluorescent signal for tumor-injected zebrafish was calculated by summing all regional scans greater than the signal cutoff for each individual sample.

All injected zebrafish displaying RFP fluorescence at 3 days post injection were selected and washed several times with clean E3 medium. Tricaine (working concentration: 0.612 nM) was then added, and the zebrafish were individually transferred to a well of a black 96-well plate for further analysis. Each zebrafish was positioned to make sure its head was centered within the well under microscope. The plate was then loaded into the TECAN microplate reader to measure RFP fluorescence intensity (excitation 555 nm, emission 585 nm). The TECAN microplate reader was configured to capture multiple data points per eachwell to ensure comprehensive coverage of the entire well area (Figure 5B), see White et al., for full methodological details.[30]

Briefly, Non-injected zebrafish were used to establish a fluorescent signal cutoff value defined as the average of the maximal fluorescence intensity value for eachnon-injected zebrafish plus three standard deviations (avg + 3SD; Figure 5C). Total fluorescent signal for tumor-injected zebrafish was calculated by summing all regional scans greater than the signal cutoff for each individual sample (Figure 5C). Fluorescence readings below the signal cutoff were considered regions of the well that did not contain a fish and thus excluded from the analysis. Fluorescence intensity was measured daily for each injected zebrafish to track changes over time. Figure 6B illustrates the relative fluorescence changes in zebrafish from Day 4 post injection, while shows fluorescence intensity persists over time.

**Figure 6:**
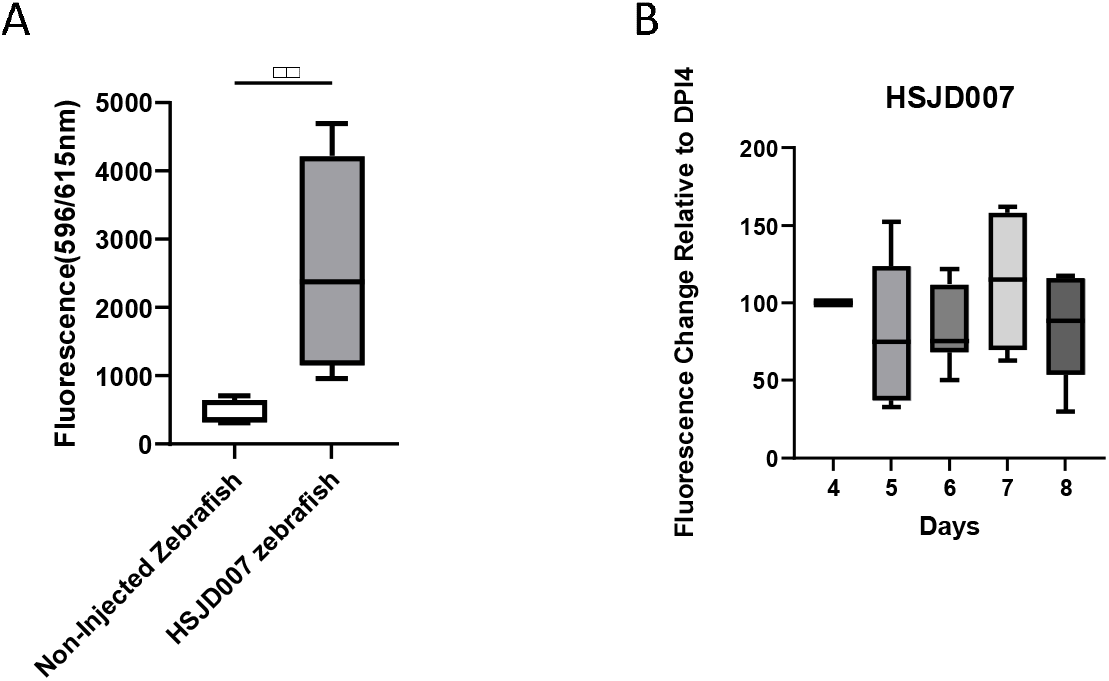
Fluorescence analysis of zebrafish injected with RFP-labeled cancer cells **A)** Comparison of fluorescence intensity between non-injected and those injected zebrafish with HSJD-007 cells. **B)** Mean fluorescence intensity change of RFP-labeled HSJD-007 cells in zebrafish over time, measured relative to 4 days post-injection (4 dpi) and monitored through 8 dpi.

### 4.7 Confirmation of DMG Cells in the brain in Zebrafish

Although time-course imaging indicated that DMG cells at the 1000-cell stage injected into zebrafish embryos migrated toward the head, it remained unclear if these cells specifically migrated into the brain. To investigate this, DMG cells were injected into transgenic zebrafish embryos expressing YFP-tagged leptomeningeal cells or microglia/macrophages (see Methods for transgene details). At six days post-injection (dpi), confocal live imaging was performed to access DMG localization.

Preparation for the whole fish imaging: Several hours before imaging, we dissolved low melt agarose (Fisher, BP1360–100) in PTU/E3 medium to achieve a final concentration of 1% and maintained it at 40°C. At least 30 minutes before imaging, we prepared the mounting solution by combining 1% LMA with tricaine, mixed thoroughly and returned the solution to 40°C. We anesthetized the fish by immersing them in Tricaine/PTU/E3, and waited approximately 3 minutes for them to become unresponsive to touch. Subsequently, we transfered individual fish into mounting medium and then positioned them within a drop (25-30 µL) of the medium on a 10 cm petri dish for imaging using an upright confocal microscope. A homemade hair loop was used to position the fish. When the agarose became solidified, the plate was moved to the confocal stage. Then, 50 ml of Tricaine/PTU/E3was added to the petri dish for imaging using water immersion objectives. Z-stacks were obtained at a 5 μm z-depth resolution.[31]

Next, “.Oib” files for each image were loaded into IMARIS (v9.7.0; Bitplane) for 3D rendering. Figure 7C demonstrates colocalization between YFP-labeled leptomeningeal cells and HSJD-007 cells, confirming the migration of DMG cells injected at the 1000-cell stage into the zebrafish brain. Figure 7D illustrates the colocalization of DIPG-2JA (RFP) cells with YFP-labeled microglia cells, showing interactions between DMG cells and the microenvironment.

**Figure 7:**
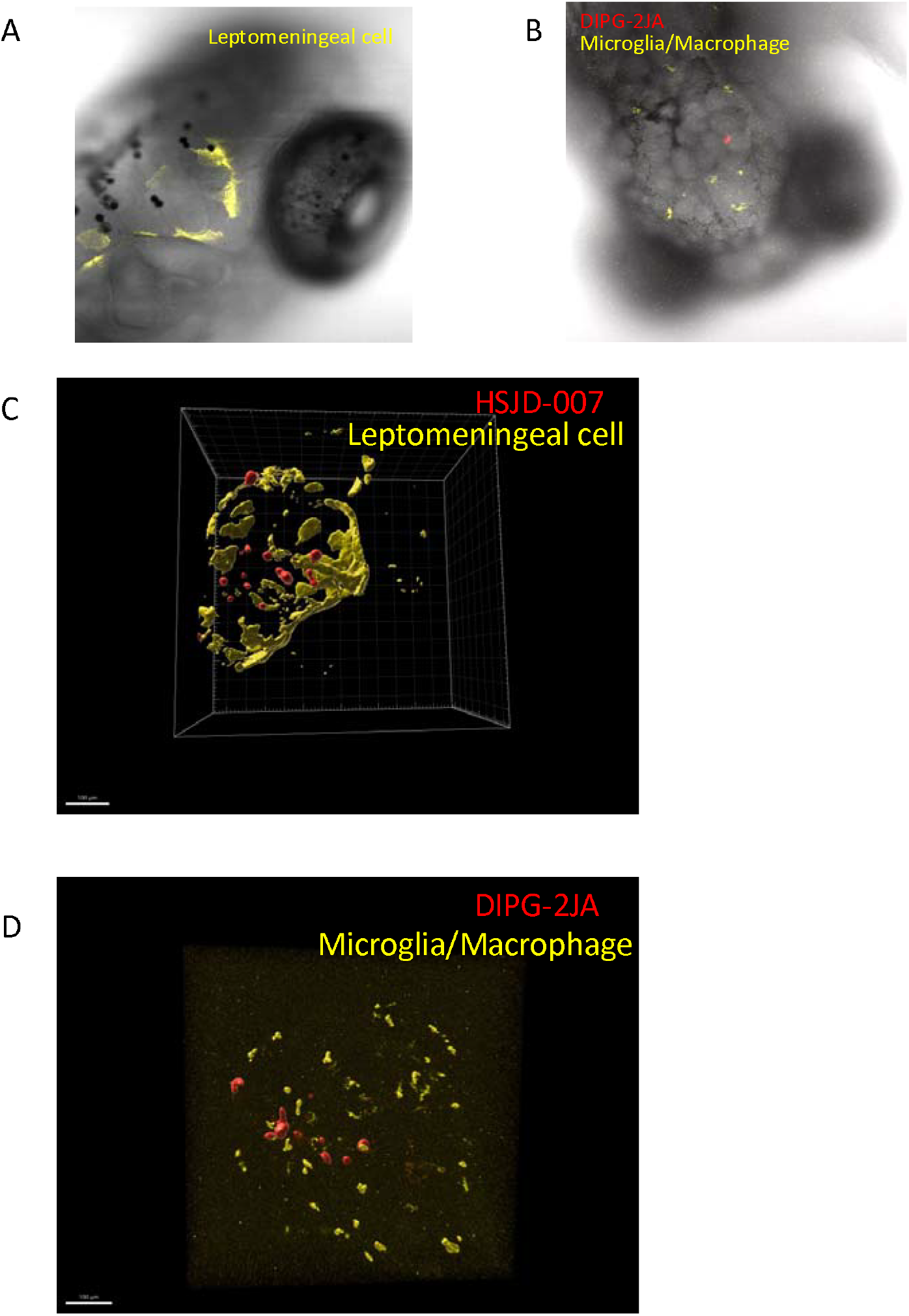
Colocalization analysis of YFP-labeled brain leptomeningeal cells and microglial cells with RFP-labeled DMG cells. **A)** Confocal image of a zebrafish head showing the position of yellow fluorescent protein (YFP)-labeled leptomeningeal cells. **B)** Confocal image of a zebrafish head displaying the position of YFP-labeled microglia and DIPG-2JA cells **C)** Colocalization of YFP-labeled brain leptomeningeal cell network (yellow) and RFP-labeled HSJD-007 tumor cells (red) in zebrafish at 6 days post-injection (dpi). **D)** Colocalization of DIPG-2JA tumor cells (red) and microglia (yellow) in zebrafish larvae at 6 days post-injection (dpi). Confocal fluorescence imaging reveals the presence of DMG cells within the brain parenchyma and spatial proximity between tumor cells and microglia/macrophages.

## 5. Conclusion

The procedure outlined here provides a robust zebrafish xenograft model for diffuse midline glioma (DMG) that is amenable to large-scale screening. The zebrafish model offers several key advantages over traditional models: rapid development, lower cost, and optical transparency, making it an efficient platform for high-throughput drug screening. By monitoring the migration of fluorescence protein positive DMG cells, researchers can track tumor burden in real time in a vertebrate model. The use of confocal microscopy enables the acquisition of time-course z-stack images, facilitating detailed assessments of tumor cell migration, proliferation, and interactions with the microenvironment. This approach significantly reduces the time and financial investment required for preclinical testing of new drugs that affect DMG viability, mobility, or the ability of DMG cells to interact with the immune microenvironment.

This model thus serves as a valuable tool for both high-throughput drug discovery and fundamental research on tumor biology. In summary, our zebrafish DMG xenograft model offers a cost-effective, scalable, and versatile platform for preclinical drug evaluation, potentially accelerating the identification of effective treatments before advancing them to mouse studies and human clinical trials.

## Acknowledgements

Funding for this research was provided by The Cure Starts Now Foundation, The Giant Food Pediatric Cancer Research Fund, and the NCI Core Grant to the Johns Hopkins Sidney Kimmel Comprehensive Cancer Center (P30CA006973).

